# Qtlizer: comprehensive QTL annotation of GWAS results

**DOI:** 10.1101/495903

**Authors:** Matthias Munz, Inken Wohlers, Eric Simon, Tobias Reinberger, Hauke Busch, Arne S. Schaefer, Jeanette Erdmann

## Abstract

Exploration of genetic variant-to-gene relationships by quantitative trait loci such as expression QTLs is a frequently used tool in genome-wide association studies. However, the wide range of public QTL databases and the lack of batch annotation features complicate a comprehensive annotation of GWAS results. In this work, we introduce the tool “Qtlizer” for annotating lists of variants in human with associated changes in gene expression and protein abundance using an integrated database of published QTLs. Features include incorporation of variants in linkage disequilibrium and reverse search by gene names. Analyzing the database for base pair distances between best significant eQTLs and their affected genes suggests that the commonly used *cis*-distance limit of 1,000,000 base pairs might be too restrictive, implicating a substantial amount of wrongly and yet undetected eQTLs. We also ranked genes with respect to the maximum number of tissue-specific eQTL studies in which a most significant eQTL signal was consistent. For the top 100 genes we observed the strongest enrichment with housekeeping genes (P=2×10^−6^) and with the 10% highest expressed genes (P=0.005) after grouping eQTLs by r^2^>0.95, underlining the relevance of LD information in eQTL analyses. Qtlizer can be accessed via http://genehopper.de/qtlizer or by using the respective Bioconductor R-package (DOI: <u>10.18129/B9.bioc.Qtlizer</u>).

## INTRODUCTION

In the past decade, genome-wide association studies (GWAS) led to the discovery of tens of thousands of associations of genetic loci in human with variation in traits and diseases. According to the NHGRI-EBI Catalog of published GWAS (GWAS Catalog), associated variants are primarily located in non-coding regions suggesting a regulatory role by modulating steps along the gene expression cascade ^1^. The most intuitive way to elucidate functional relationships between regulatory variants and genes is to investigate the correlation between variant occurrence and gene expression. The resulting data type is referred to as expression quantitative trait locus (eQTL) if a variant (i.e. the QTL) influences mRNA levels or protein quantitative trait loci (pQTL) if a variant influences a protein level.

Along with the growing number of associations from GWAS, many QTL databases (DBs) have evolved: the GTEx Portal ^2^, Haploreg ^3^, GRASP ^4^, GEUVADIS ^5^, SCAN ^6^, seeQTL ^7^, Blood eQTL Browser ^8^, pGWAS ^9^, ExSNP ^10^ and BRAINEAC ^11^. Some of them, but in particular the GTEx Portal, provide specific tools and visualizations to analyze their QTL data. However, most of the existing DBs only provide unique QTL datasets resulting in a poor overlap between the DBs. In addition, the databases are very limited regarding the available search term types and do not permit searching for multiple variants or genes in parallel. Hence, it is very cumbersome to comprehensively annotate lists of variants or genes with immediate results. Another aspect that the current databases do not sufficiently consider, is that for both GWAS and QTL studies, true association signals are very often accompanied by multiple other variants, which can be attributed to the linkage disequilibrium (LD) structure in the human population. The correlation between variants at a locus due to LD makes it difficult to identify causal variant(s) just as it is hard to find true signal overlaps between a GWAS and a QTL signal.

To address these shortcomings, we have developed the web-based application “Qtlizer”, which facilitates and improves QTL annotation. Qtlizer allows exploration of QTL data for a given list of small variants (Indels and SNPs) and/or genes in a fast and efficient manner by integrating a large number of QTL datasets. All data is extensively annotated in the web application and direct links to the source platform are provided for further analyses. To enhance the interpretability regarding signal overlaps, Qtlizer allows for the inclusion of variants in LD with a query variant, as well as flag them, which helps to dissect the QTL data into independent association signals. In line with the release of Qtlizer, we provide two unique eQTL datasets from the Cardiogenics Consortium ^12,13^.

Apart from the aforementioned existing QTL platforms many other tools exist for annotating and characterizing variants resulting from GWAS studies, for example FUMA ^14^ and SNiPA ^15^. These tools, however, target a more general annotation of variants and are restricted to a small number of eQTL datasets. Therefore, we did not compare them with Qtlizer in more detail in this work.

## MATERIALS AND METHODS

### Data sources

Qtlizer currently integrates 167 tissue-specific QTL studies of which two are response expression QTLs (reQTLs) and pQTLs, while the remainder are eQTLs. Further details are provided in **Supplementary Table 1**. eQTL data was obtained from 13 sources that included datasets from 53 publications and cover 114 different tissues or cell types. For 34 of these tissues more than one eQTL dataset is available, with most included studies performed for lymphoblastoid cell lines (n=9), liver (n=8), cerebellum (n=4) and monocytes (n=4). The number of variants per study varies considerably and reflects the diversity of the included data; the median is approx. 3,700, but the number of variants ranges up to 2,017,804 for a thyroid dataset from the GTEx Portal. Some of the included datasets cover only variants that are statistically significant after multiple testing in the original study, while others also cover suggestively associated SNPs.

Moreover, we included several other datasets. Published variant phenotype associations identified in GWAS were taken from GWAS Catalog ^16,17^. We used genotype data from 1000 Genomes Project Phase 3 (1000GP3) ^18^. A dataset of topological associated domain (TAD) boundaries was taken from Dixon et al. ^19^. TADs are spatially close genetic regions in the genome and have been defined based on Hi-C chromatin interaction data, i.e. physical interactions occur more frequently within such a compartment. All other data such as genetic variant and gene information was taken from the Genehopper DB (http://genehopper.de) ^20^. Genehopper DB is an integrated DB that mainly relies on data collected from Ensembl ^21,22^.

### Web server implementation

We used the programming language Perl to implement an extraction, transformation and loading (ETL) process in which QTL summary statistics and other aforementioned data were downloaded from the respective websites (extraction), manipulated to fit the database schema of the host database (transformation) and the resulting tables were loaded into the host database (**Figure 1A, B**). As a host database we used the Genehopper DB which is based on the relational database management system MySQL (https://www.mysql.com). The DB was then connected to the web application which we built using the Play! Framework (https://www.playframework.com) (**Figure 1C**). We used Java for implementing the backend and Javascript for implementing the frontend of the application. Lastly, we used the front-end component library Bootstrap (https://getbootstrap.com/docs/3.3/getting-started/) to give Qtlizer a responsive and appealing design.

**Figure 1.**
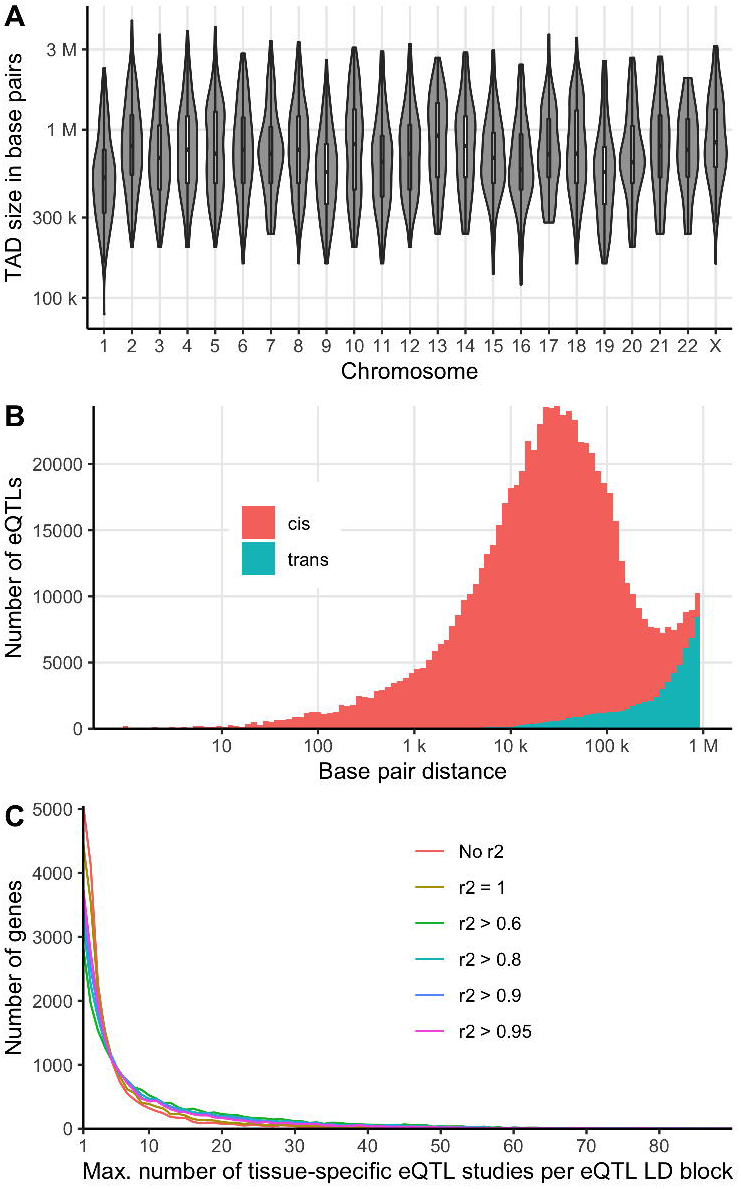
**A** Data from various publicly available resources was integrated **B** applying an Extraction, Transformation and Loading (ETL) process. **C** Qtlizer consists of a relational database and a web application **D** Qtlizer can be queried for QTL data via the web-based user interface or programmatically by inputting lists of genetic variants and genes using a REST API. **E** Annotation results are displayed in a table view.

## RESULTS

### Data transformation

After selecting and downloading the datasets described in the previous section, we applied a transformation procedure as part of the ETL process. This procedure consisted of several crucial steps. We manually curated tissue names to make them consistent across all datasets. If available, we took QTL significance information from the source. Otherwise, the significance was determined by adjusting for multiple testing using a family-wise error rate (FWER) of 5%, resulting in an adjusted significance level of P = 10^−12^ which corresponds to 1000,000 variants and 50,000 genes (approximately equals to the set of Ensembl genes without pseudogenes). eQTL objects that passed the study-wide significance threshold were flagged as “is_sw_significant”. Here and in the following we define a QTL object as a quadruple of variant, gene, tissue and source study. We utilized the TAD boundaries to categorize QTLs into *cis* or *trans* which denotes whether a QTL gene-variant pair is located within the same TAD or not. All variants and genes, that were included in the QTL data, were mapped to the GWAS Catalog for possible variant-phenotype or locus-phenotype associations. Variant-variant correlations were calculated as LD based on the 1000GP3 genotypes of the European population **(Supplementary Figure 1)**. For each QTL object we used this LD information to investigate whether the underlying variant had the lowest P-value among all variants within its LD group in a specific tissue of a QTL study and flagged the QTL object accordingly with “is_best_in_ld_group”. We defined the LD group of a variant as the variant and all other variants with r^2^ > 0.2. Moreover, a QTL object was flagged as “is_best” if it had the overall lowest P-value in a specific tissue of a QTL study. Lastly, we calculated count characteristics for each QTL. For each gene in a specific tissue of a QTL study, we calculated (i) the number of putative QTLs (“n_qtls”), (ii) the number of tissues and studies in which the putative QTL is best with respect to the P-value (“n_best”), (iii) the number of tissues and studies for which the putative QTL has the best study-wide significance (“n_sw_significant”) and (iv) the number of tissues and studies in which the putative QTL occurs (“n_occ”).

40,883,209 (37,014,094 study-wide significant) QTLs from 3,856,968 variants and 32,987 genes were finally added to the Genehopper DB. Of these, 25,432,954 QTLs were annotated as being *cis* QTLs and 26,449 variants and 11,511 genes could be linked to an entry in the GWAS Catalog. We integrated boundaries of 3,054 TADs with a mean length of 853 kilo bases (kb; maximum length = 4.44 mega bases, minimum length = 0.8 kb) (**Figure 2A**).

**Figure 2.**
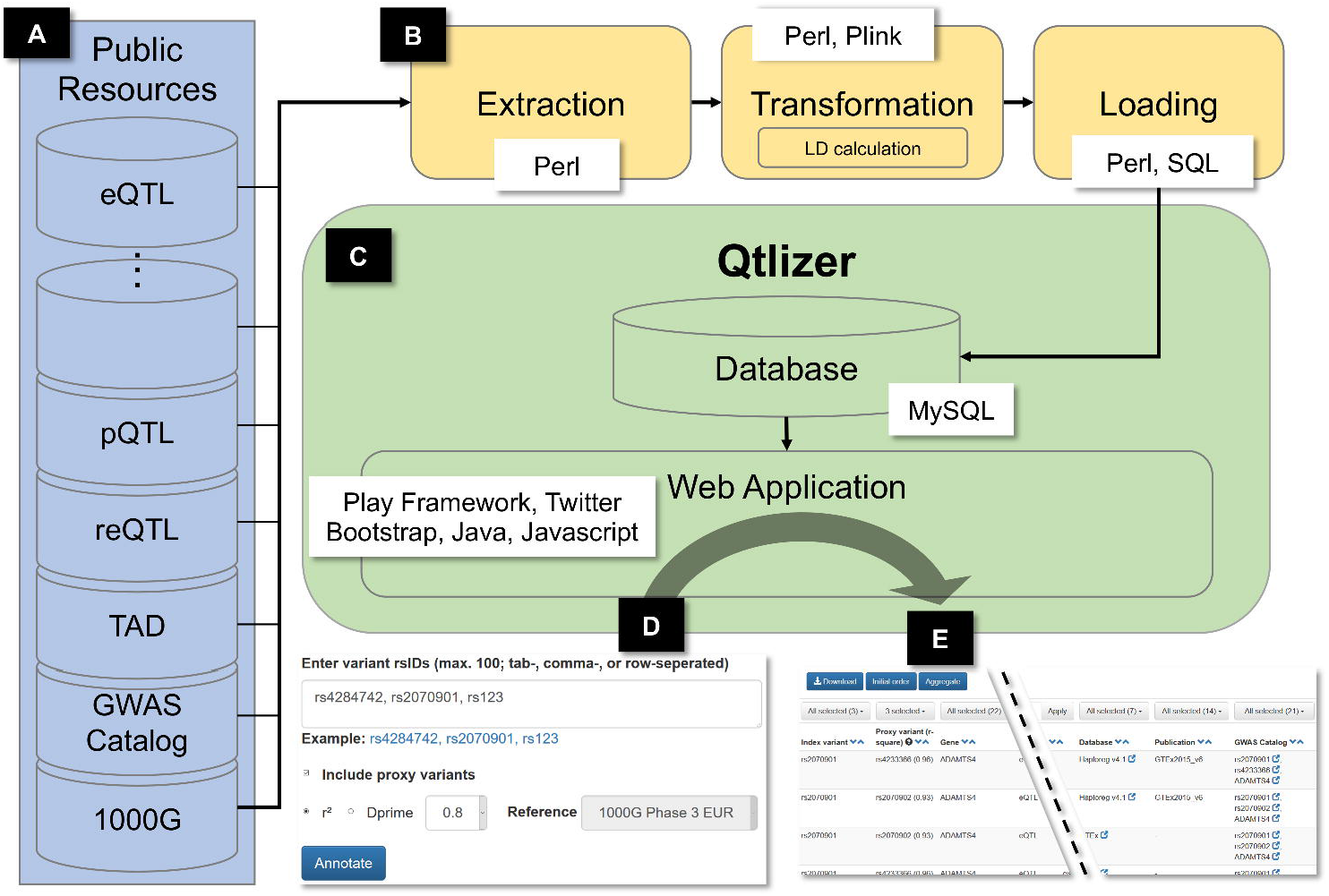
**A** Per chromosome distribution of the TAD sizes in base pairs **B** Distribution of the base pair distances between best eQTLs and affected genes in the GTEx dataset. **C** Distribution of the maximum number of tissue-specific eQTL studies per eQTL LD block for each gene.

## The web application

### User interface workflow

The user interface of Qtlizer takes a list of query terms as input (**Figure 1D**). A single query term can identify either a genetic variant or a gene. Accepted variant identifiers are reference SNP identifiers (rsIDs) or chromosomal positions of the reference builds hg19/GRCh37 or hg38/GRCh38. Genes can be specified using a variety of database identifiers (e.g. Ensembl Gene Id, EntrezGene Id) or gene symbols consisting of upper-case letters and Arabic numerals. Gene terms are mapped to an internal gene identifier by using the full text term-to-gene index and a prioritization algorithm provided by Genehopper to account for ambiguous identifiers or symbols^20^. By using regular expressions, Qtlizer requires no special input format and is able to interpret all standard text formats (e.g. tab-, comma-, space-separated, or a mixture of all). The regular expressions are listed in **Supplementary Figure 2**.

Optionally, queried variants can be enriched for proxy variants using the European reference population of 1000GP3 by setting a LD threshold for r^2^ or D’.

The query result is displayed in a table view, with one row for every QTL object (**Figure 1E**). This means that each row represents a putative association of a variant with the expression of a gene in a given tissue and within a QTL study that has been obtained from the given source.

Initially, only elementary information is displayed: (i) variant rsID and hyperlinks to dbSNP and Ensembl; (ii) rsID of the proxy variant, if input variants were enriched for proxies; (iii) gene symbol with a hyperlink to Ensembl; (iv) chromosomal distance in kilo bases between variant and gene; (v) normalized tissue name; (vi) association P-value; (vii) beta value representing the effect size; (viii) effect and non-effect alleles (EA, NEA); (ix) source name with hyperlinks to the source database and original publication in which the dataset was published; (x) flags “is_significant”, “is_best”, “is_best_in_ld_group” which are grouped in a single column; (xi) hyperlinks that connect the variant, proxy variant and/or the gene with entries in the GWAS Catalog.

Additionally, there are a number of optional columns that can be enabled. The search type can be a “Variant” if the query term was mapped to a variant, or “Gene” if the query term was mapped to a gene; the original query term that was entered by the user; the type of QTL, e.g. eQTL or pQTL; Ensembl Gene ID; co-localization (*cis* versus *trans*); a significance information column providing details about the multiple testing correction and the applied significance threshold; the Pubmed ID of the original publication and a counts column summarizing the data for several eQTL studies.

The resulting table can be interactively sorted and filtered column-wise, either by selecting an arbitrary subset of column values (e.g. a selection of SNPs or genes or tissues) or by specifying numerical thresholds (e.g. P-value < 0.001 or beta > 0). The current filter state of the table can be downloaded as text file for further use.

Qtlizer allows summarizing the results such that for each index variant and gene all available tissues and the best P-value are merged into a single row. QTLs of proxy variants are merged with the index variant to allow for a haplotype block-centric view.

### Example use case

We show how to use Qtlizer for annotating a coronary artery disease (CAD) GWAS locus at chromosome 3q22.3, whose index SNP is rs2306374 (**Supplementary Figure 3**) ^23^. After specifying the input SNP and clicking the button “Annotate”, Qtlizer lists 31 QTL records (indicated as “#Results: 31”) for this SNP in the Qtlizer database. After enabling the column “QTL type” using the button “Select columns”, it turns out that all records are eQTLs. By clicking the button “Aggregate”, we see that there are seven eQTL genes and that SNP rs2306374 is associated with the expression of *MRAS* (Muscle RAS oncogene homolog) in multiple tissues. Most of these tissues are relevant for coronary artery disease, e.g. “Artery – Aorta”, “Artery – Tibial”, “Heart - Atrial appendage” and “Heart”. By clicking on the SNP in the “GWAS Catalog” column, we learn from the GWAS Catalog website that the SNP’s C allele is associated with CAD according to a GWAS performed in altogether more than 100,000 Europeans ^24^. After switching back to Qtlizer’s non-aggregated table view by clicking the “Aggregate” button again, we investigate the *MRAS* gene closer by selecting it exclusively in the “Gene” column header. 14 records remain, all being study-wide significant, i.e. flagged as “is_sw_significant” which denotes they are study-wide significant eQTLs. By inspecting the allele columns “EA” and “NEA”, we observe that the eQTL effect allele (EA) is C and from the “Beta” column we see that in six CAD-relevant tissues *MRAS* expression is increased for the EA, whereas it is decreased in six other tissues. The “Distance” column specifies a distance of 0 which denotes that SNP rs2306374 is located within the gene boundaries. The absence of the “is_best” and “is_best_in_ld_group” flags denotes that the query SNP is not the eQTL sentinel variant.

In a next step we start a new Qtlizer search which includes proxy variants. We choose a conservative LD threshold of r^2^ = 0.8, resulting in 585 associations. The aggregate view confirms that *MRAS* is still the eQTL gene observed in most relevant tissues, and additional relevant tissues for proxies are “Atherosclerotic aortic root” and “Artery – Mammary”. We again select only *MRAS* eQTLs and filter further to keep those associations in heart, artery/aortic tissues tagged with flags “is_sw_significant” and one or both of “is_best”. We obtain 11 associations for five proxy variants with r^2^ = 0.94 (provided in brackets behind the respective rsIDs). Two sources, GTEx and Haploreg, concordantly report the eQTL signal as being best in four heart tissues, and two sources, GTEx and Franzén et al., as being best for the LD block in five artery tissues.

Sorting the records by beta which denotes the effect size, we select SNP rs13324341 (r^2^ = 0.94; beta = 1.04) for functional studies. SNP rs13324341 is intronic of *MRAS* and lies in a DNase I hypersensitivity site according to the ENCODE track in the UCSC Browser ^25,26^. Moreover, according to the “Count” column, this SNP is the best QTL for *MRAS* in four studies in artery tissues.

## Explorative analyses

### Distance between genes and eQTLs

We used our integrated database to study the base pair distance distribution between eQTL and gene per tissue/source pair by calculating the distance from gene start position to the position of the best (regarding P-value) study-wide significant eQTL**. Figure 2B** shows the log scaled distance distributions across all tissues in the GTEx dataset. The left tail of the *cis* curve appears much longer and thicker than the right tail, while the *trans* curve (and total) are skewed the other way. For the total distribution we measured a mean distance of 101 kilobase pairs (kb) and a median distance of 28 kb. The sharp cutoff at 1 Mb is due to the *cis*-distance limit for variant and gene comparisons that was set by GTEx and that is commonly used for eQTL calculations ^27^. Accordingly, the distances in the other datasets showed a similar distribution pattern. Intuitively, when extrapolating the distribution towards distances with more than 1 Mb, a less conservative cutoff might reveal a substantial number of as-yet unseen eQTLs.

### Stable eQTLs across multiple tissues

Next, we studied the robustness of eQTL signals across different studies, i.e. in how many tissue-specific eQTL studies (i.e. source/tissue pairs) the same eQTL was found for a specific gene. Since only few source publications refer to the same tissue (e.g. liver), this assessment very well approximates the sharing across tissues. Again, we only considered the best study-wide significant eQTL per tissue-specific eQTL study and gene. For each gene, eQTLs were grouped into LD blocks (in the following referred to as eQTL LD block) for different r^2^ thresholds (r^2^ > 0.6; r^2^ > 0.8, r^2^ > 0.9; r^2^ > 0.95; r^2^ = 1; No LD). Subsequently, we counted the number of tissue-specific eQTL studies for each eQTL LD block and assigned the eQTL LD block with the maximum number of tissue-specific eQTL studies to each gene. 18,541 genes had at least one best study-wide significant eQTL. For all thresholds, we found that the number of genes exponentially decreases with the number of tissue-specific eQTL studies per eQTL LD block (**Figure 2C**). Our findings support the tissue-by-tissue sharing pattern of genes associated with expression as reported by GTEx under inclusion of more than twice as many tissues ^28^. We confirm that expression of a significant proportion of genes is genetically controlled in a single tissue or a small subset of tissues. Additionally, by considering only the best eQTL per gene and tissue-specific study and by identifying the LD block with maximum number of such eQTLs, we find that expression is often strongly associated with the same set of correlated genetic variants in a small number of tissues (see **Figure 2C**). Thus, genetic control likely acts in multiple tissues via the same causal variant(s) and accordingly possibly via the same underlying biological mechanism.

Gene *FAM118A* (Family with sequence similarity 118 member A) showed the highest count of tissue-specific eQTL studies (n > 100) per eQTL LD block for r^2^ thresholds < 1. For each r^2^ threshold, we investigated the 100 genes with the highest maximum count of tissue-specific eQTL studies for overlap with housekeeping genes compiled by Eisenberg and Levanon ^29^ and with the top 10 % highest expressed genes (mean expression across all tissues) in GTEx (**Supplementary Table 2**). We observed the strongest enrichment with housekeeping genes at a grouping threshold r^2^ > 0.95 (False discovery rate < 0.05 adjusted P = 2×10^−6^, hypergeometric test) (**Supplementary Table 3**). Similarly, we observed the strongest enrichment with highly expressed genes at the same threshold (Adjusted P = 0.03), whereas no enrichment was detected without grouping (Adjusted P = 0.2).

## DISCUSSION

With Qtlizer we introduce a web-based tool to annotate lists of common small variants and genes with QTL data in a comprehensive way. Qtlizer has an easy-to-use graphical interface for jointly reviewing many QTL studies on various tissues, covering a large portion of human common small variants and genes. This is possible because we integrated QTL data from many different sources, connected them with various other data types and calculated meaningful characteristics. To-date, Qtlizer contains the most diverse and comprehensive set of QTL data. Moreover, it provides unique functionality to analyze multiple inputs in parallel.

It is important to note that we explicitly did not aim to statistically combine the QTL data over studies or tissues. Such efforts are difficult because studies differ in various aspects such as sample sizes, disease status and genetic background of genotyped individuals as well as techniques used for genotyping and expression quantification (e.g. array versus sequencing). Besides these differences, there are also other restrictions regarding the public availability. For example, some studies provide only significant associations after multiple testing correction whereas others use thresholds on nominal association P-values. Another challenge is missing information. For example, beta and effect alleles are occasionally not reported and as a result the effect direction cannot be determined. Due to QTL data heterogeneity, Qtlizer is designed to provide comprehensive access to as many available QTL datasets as possible in order to provide a starting point for in-depth literature review and further analysis in native QTL browsers, if available.

Also, we refrained from filtering variants or genes to be included in Qtlizer to enable the user to review all available information. However, we do allow interactive prioritization of variant-gene relationships.

In genetic association studies such as GWAS and QTL studies, association signals are strongly influenced by the LD structure. This means that a true association signal usually does not only consist of the causal variant(s) but is supported by variants which are correlated with the causal variant(s), producing an inherent ambiguity in interpreting association study results. This phenomenon is visualized as coherent elevations in regional association plots for which we provide an example in **Supplementary Figure 3**. Consequently, the problem of correlated variants makes it also difficult to determine whether two association signals overlap and point to the same causal variant(s). A typical case use for Qtlizer is to compare a GWAS signal, which is represented by an index variant given as input by the user, with that of a QTL signal in Qtlizer. To assist the user with investigating the signal overlap between the query and the QTL data in Qtlizer, we implemented the option to include LD variants and to prioritize the QTLs using informative flags and summary counts. Flags that we want to highlight in this context are “is_best” and “is_best_in_ld_group”, informing the user whether a QTL object is best among all correlated QTLs in a tissue and study regarding the association P-value. If sufficient data is available for the tissue of interest, the user can also export selected QTL data for testing the significance of overlap of association signals using dedicated tools such as Coloc ^30^.

It is common practice to divide between *cis* eQTLs and *trans* eQTLs. Usually, the functional mechanism is not known and eQTL studies use the term *cis*, if the variant and the associated gene are close to each other (e.g. within 1 MB) and *trans* otherwise. In Qtlizer we improve on this crude definition by using TADs boundaries. Although multiple publications have suggested that chromosomal TAD boundaries are largely conserved across cell-types ^19,31,32^, recent studies indicate that TAD boundaries might be more unstable than previously assumed ^33^. However, in this work, we assumed the former. High quality, tissue-specific TAD boundaries can be included in Qtlizer once available.

The frontend functionality of Qtlizer was implemented in Javascript which means, that the performance of rendering and manipulating the result table is highly dependent on the hardware resources of the user. For this reason, we restricted queries to a maximum of 50 query terms and a maximum table size of 10,000 entries. For queries exceeding these thresholds, we allow programmatic access to the QTL data by using our Bioconductor R-package or by using our web service directly. For the latter, we provide example scripts for Perl and R in **Supplementary Figure 4** and **Supplementary Figure 5**.

Keeping the database of Qtlizer up to date is especially important when the source databases release new versions or new datasets are being published. Therefore, we implemented a semi-automated ETL process allowing to update Qtlizer in a timely and simple manner.

Apart from the data integration and web application efforts we also explored our integrated QTL database for new insights. Investigating the distances between best eQTLs and their affected genes suggests that the commonly used 1 Mb distance threshold as upper bound for describing variant-gene relationships in *cis* might be too restrictive for detecting all best eQTLs and lead to wrong assignments. Our observation is also supported by fact that the TAD boundaries we utilized which can span multiple millions base pairs. We also ranked genes according to the number of tissue-specific eQTL studies in which a single affecting eQTL was most strongly associated with expression. As expected, the top genes were enriched for highly and constitutively expressed genes. Interestingly, this result was most obvious after grouping eQTLs into correlated groups using LD information. This observation highlights the importance of including proxy variants in the eQTL analyses.

In summary, Qtlizer provides a high performant web-based solution for annotating lists of genetic variants and genes with QTLs, facilitating the exploration of the consequential genetic associations in non-coding regions on the molecular mechanisms of a phenotype. Qtlizer outperforms each of the existing databases regarding the number of QTL datasets and the variety of tissues that can be explored. Since QTLs can be tissue-specific, this is particularly helpful to select appropriate tissues for a phenotype of interest. In addition, we have implemented an ETL pipeline that facilitates the integration of new QTL datasets to keep Qtlizer updated.

## Supporting information

Supplementary Material (Excel Sheet)

## AVAILABILITY

The web application of Qtlizer is available at http://www.genehopper.de/qtlizer. The website is free and open to all and there is no login requirement. Qtlizer can also be accessed in R by using the respective Bioconductor package (DOI: <u>10.18129/B9.bioc.Qtlizer</u>). A guide on how to use the REST API directly is available at http://www.genehopper.de/rest (**Supplementary Figure 4, 5**).

## ACKNOWLEDGMENTS

We acknowledge computational support from the OMICS compute cluster at the University of Lübeck. Special thanks to Dr. Carsten Kemena for extensive testing of the web application, Dr. Loreto Munoz Venegas for revising the manuscript, Julia Remes who contributed to the R package of Qtlizer and to Lena Friedrichsen for her support in the variant-gene distance analyses. We cordially thank the Cardiogenics Consortium for data related to the macrophage and monocyte eQTLs. Members of the Cardiogenics Consortium are listed in **Supplementary Table 4**.

## AUTHOR CONTRIBUTIONS

M.M. developed and implemented Qtlizer and wrote the first draft of the manuscript. I.W. wrote the documentation and contributed to the manuscript. E.S., H.B., A.S., J.E., T.R. contributed to the manuscript and tested Qtlizer.

