## Supplementary Material for "Qtlizer: comprehensive QTL annotation of GWAS results"

**Supplementary Figure 1.** Linkage disequilibrium (LD) was calculated with PLINK 1.9 beta (https://www.cog-genomics.org/plink2/). The script shows the parameters that we passed to PLINK for calculating LD. Among these, `--ld-window` sets the maximum window size as number of variants, whereas `--ld-window-kb` sets the maximum window size in kilo bases (kb). PLINK considers the window size which is reached first. Given a variant, the window size determines for how many downstream (regarding chromosomal position) variants the LD should be calculated for a given variant.. 4
**Supplementary Figure 2.** Regular expressions that are used by Qtlizer to extract (a) rsIDs, (b) chromosomal positions and (c) gene identifier. These expressions are applied in the backend (Java) and frontend (Javascript) of Qtlizer. Regarding Javascript interpretation, it has to be stated that, to-date, lookbehind assertions `?<=` are only be supported by the Google Chrome browser, but not by Firefox, Safari and Edge. Therefore, we converted the regexes accordingly (d-f)...................................................... 5
**Supplementary Figure 3.** Regional association plot showing an association signal on chromosome 3 at the gene locus MRAS. Each dot represents a single nucleotide polymorphism (SNP). The colour coding reflects the degree of linkage disequilibrium ($r^2$) between a variant and the index variant rs2306374. This plot visualizes GWAS data on coronary artery disease provided by the CARDIoGRAMplusC4D Consortium (http://www.cardiogramplusc4d.org/data-downloads/). ......................................................... 6
**Supplementary Table 3.** For each LD threshold the number of housekeeping genes and the number of genes in the 10% highest expressed genes in GTEx v8 among the 100 genes with the highest dataset count per eQTL LD block. Adjusted (FDR < 0.05) P-values were calculated using hypergeometric test assuming an overall gene number of 18,521 (genes with at least one best significant eQTL), a total of 3,705 housekeeping genes and a total of 1,201 genes among the 10% highest expressed genes. .............. 13
**Supplementary Table 4.** Members of the Cardiogenics Consortium. ..................................................... 14

```
#!/bin/bash

/path/to/plink --bfile <bim/bed/fam prefix> --r2 --ld-window 99999 --ld-window-kb
1000 --ld-window-r2 0.2 --out <output_file>
```

**Supplementary Figure 1.** Linkage disequilibrium (LD) was calculated with PLINK 1.9 beta (https://www.cog-genomics.org/plink2/). The script shows the parameters that we passed to PLINK for calculating LD. Among these, `--ld-window` sets the maximum window size as number of variants, whereas `--ld-window-kb` sets the maximum window size in kilo bases (kb). PLINK considers the window size which is reached first. Given a variant, the window size determines for how many downstream (regarding chromosomal position) variants the LD should be calculated for a given variant.

```
a)

"(?<=^|[\\s,;])rs(\\d+){1,12}(?=[\\s,;]|$)"

b)

"(?<=^|[\\s,;])(hg19|hg38|grch37|grch38)[:_-](chr)?(1[0-9]|2[0-2]|[1-
9]|x|y|mt)[:_-](\\d+){1,8}(?=[\\s,;:_-]|$|[atgc]+)"

c)

"(?<=^|[\\s,;])([a-z0-9]+){2,20}(?=[\\s,;]|$)"

d)

Array.from(new Set(q.match(/(^|[\s,;])rs(\d+){1,12}(?=[\s,;]|$)/g)))
      .map(function(e) {return e.replace(/^[\s,;]/g, "")});

e)

Array.from(new Set(q.match(/(^|[\s,;])(hg19|hg38|grch37|grch38)[:_-](chr)?(1[0-
9]|2[0-2]|[1-9]|x|y|mt)[:_-](\d+){1,8}(?=[\s,;:_-]|$|[atgc]+)/g)))
      .map(function(e) {return e.replace(/^[\s,;]/g, "")
                                .replace(/_|-/g, ":")
                                .replace(/grch37/g, "hg19")
                                .replace(/grch38/g, "hg38")
                                .replace(/chr/g, "");});

f)

Array.from(new Set(q.match(/(^|[\s,;])([a-z0-9]+){2,20}(?=[\s,;]|$)/g)))
      .map(function(e) {return e.replace(/^[\s,;]/g, "")})
                                .filter(function(v){return !v.match(/^rs\d+$/)});
```

**Supplementary Figure 2.** Regular expressions that are used by Qtlizer to extract (a) rsIDs, (b) chromosomal positions and (c) gene identifier. These expressions are applied in the backend (Java) and frontend (Javascript) of Qtlizer. Regarding the Javascript implementation, lookbehind assertions `?<=` have only recently been supported by all common browsers. To ensure backwards compatibility, we converted the regexes accordingly (d-f).

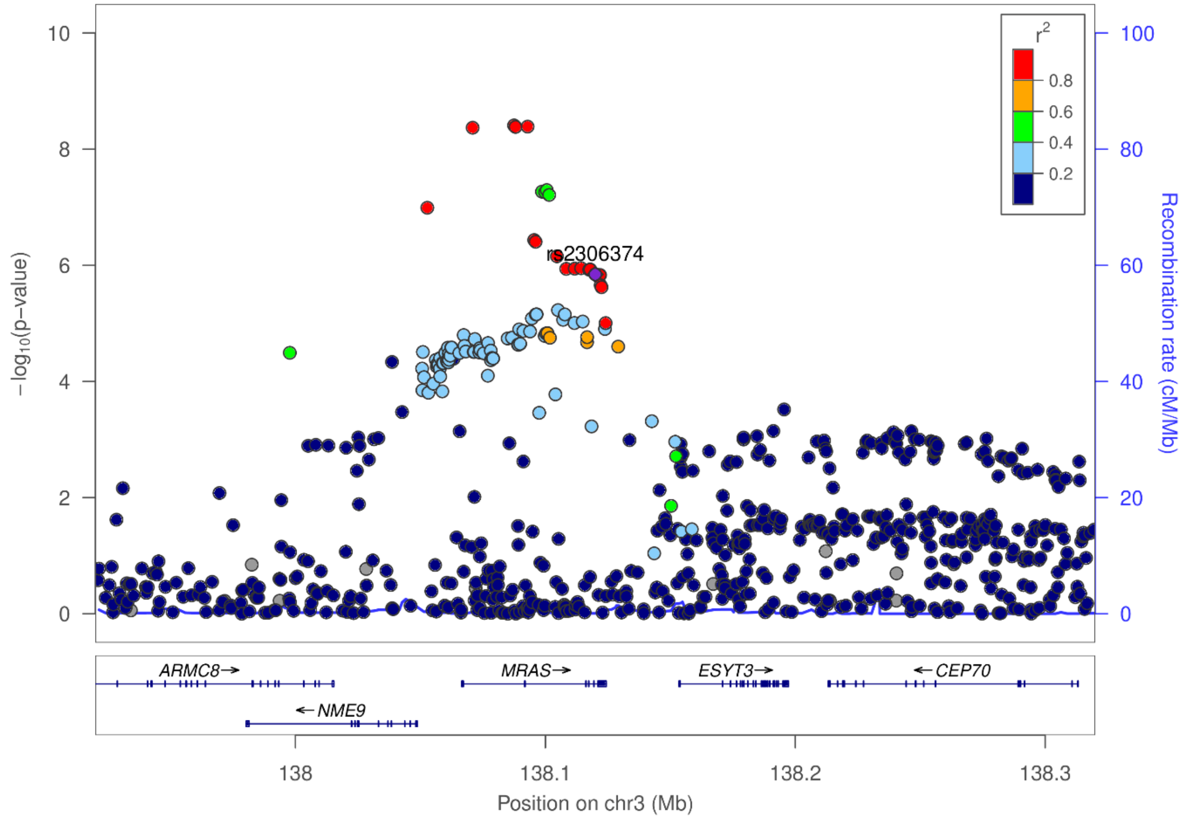

**Supplementary Figure 3.** Regional association plot showing an association signal on chromosome 3 at the gene locus *MRAS*. Each dot represents a single nucleotide polymorphism (SNP). The colour coding reflects the degree of linkage disequilibrium ($r^2$) between a variant and the index variant rs2306374. This plot visualizes GWAS data on coronary artery disease provided by the CARDIoGRAMplusC4D Consortium (http://www.cardiogramplusc4d.org/data-downloads/).

```
# https://cran.r-project.org/web/packages/httr/vignettes/quickstart.html
library(httr)

q = "rs4284742,rs2070901,rs123"
corr = 0.8 # optional
ld_method = "r2" # optional

url = paste('http://www.genehopper.de/rest/qtlizer?q=', q, "&corr=", corr,
"&ld_method=", ld_method, sep="")

response = POST(url);
result = content(response)

a = unlist(strsplit(result , "\n|\t"))
m = matrix(a[20:length(a)], ncol=19, byrow=TRUE)
d = as.data.frame(m, stringsAsFactors=FALSE)
colnames(d) = a[1:19]

write.table(d, file="example.txt", sep="\t", quote = FALSE, append=FALSE,
row.names = FALSE, col.names = TRUE)
```

**Supplementary Figure 4**. For larger queries Qtlizer can be accessed using the Genehopper REST API. The script describes an example request in **R**. More info about the API can be found at http://www.genehopper.de/rest.

```perl
#!/usr/bin/perl

use warnings;
use strict;
use LWP::UserAgent;

my $url = 'http://www.genehopper.de/rest/qtlizer';

my %parameters = (
        "q" =>  "rs4284742, rs2070901, rs123",
        "corr" => 0.8, # optional
        "ld_method" => "r2" # optional
);

my $url_with_pars = "$url?";
$url_with_pars .= $_."=".$parameters{$_}."&" for keys(%parameters);

my $ua = LWP::UserAgent->new();
my $response = $ua -> post($url_with_pars);
my $content = $response -> decoded_content();

print $content."\n";
```

**Supplementary Figure 5.** The script describes an example request in **Perl** (http://www.genehopper.de/rest).

**Supplementary Table 1.** List of QTL datasets that were integrated into Qtlizer.

| Source | PMID | Tissue | QTL Type | #Variants |
|---|---|---|---|---|
| Blood eQTL Browser | 24013639 | Peripheral blood | eQTL | 768850 |
| Franzén et al. (Science 2016) | 27540175 | Adipose - Subcutaneous | eQTL | 6776 |
| Franzén et al. (Science 2016) | 27540175 | Adipose - Visceral (Abdominal) | eQTL | 6697 |
| Franzén et al. (Science 2016) | 27540175 | Adrenal gland | eQTL | 1 |
| Franzén et al. (Science 2016) | 27540175 | Artery - Aorta | eQTL | 288 |
| Franzén et al. (Science 2016) | 27540175 | Artery - Mammary | eQTL | 7109 |
| Franzén et al. (Science 2016) | 27540175 | Artery - Tibial | eQTL | 12 |
| Franzén et al. (Science 2016) | 27540175 | Atherosclerotic aortic root | eQTL | 6958 |
| Franzén et al. (Science 2016) | 27540175 | Blood | eQTL | 5591 |
| Franzén et al. (Science 2016) | 27540175 | Brain - Hypothalamus | eQTL | 2 |
| Franzén et al. (Science 2016) | 27540175 | Brain - Nucleus accumbens (Basal ganglia) | eQTL | 2 |
| Franzén et al. (Science 2016) | 27540175 | Brain - Putamen (Basal ganglia) | eQTL | 11 |
| Franzén et al. (Science 2016) | 27540175 | Cells - Transformed fibroblasts | eQTL | 658 |
| Franzén et al. (Science 2016) | 27540175 | Colon - Transverse | eQTL | 18 |
| Franzén et al. (Science 2016) | 27540175 | Esophagus - Mucosa | eQTL | 976 |
| Franzén et al. (Science 2016) | 27540175 | Esophagus - Muscularis | eQTL | 15 |
| Franzén et al. (Science 2016) | 27540175 | Heart - Left ventricle | eQTL | 3 |
| Franzén et al. (Science 2016) | 27540175 | Liver | eQTL | 7043 |
| Franzén et al. (Science 2016) | 27540175 | Lung | eQTL | 98 |
| Franzén et al. (Science 2016) | 27540175 | Muscle skeletal | eQTL | 5650 |
| Franzén et al. (Science 2016) | 27540175 | Nerve - Tibial | eQTL | 30 |
| Franzén et al. (Science 2016) | 27540175 | Pancreas | eQTL | 283 |
| Franzén et al. (Science 2016) | 27540175 | Prostate | eQTL | 1 |
| Franzén et al. (Science 2016) | 27540175 | Skin - Not sun exposed (Suprapubic) | eQTL | 11 |
| Franzén et al. (Science 2016) | 27540175 | Skin - Sun exposed (Lower leg) | eQTL | 64 |
| Franzén et al. (Science 2016) | 27540175 | Testis | eQTL | 203 |
| Franzén et al. (Science 2016) | 27540175 | Thyroid | eQTL | 2120 |
| Franzén et al. (Science 2016) | 27540175 | Whole blood | eQTL | 35 |
| GEUVADIS | 24037378 | Cells - Lymphoblastoid cell lines | eQTL | 404608 |
| GRASP 2 Catalog | 17873877 | Cells - Lymphoblastoid cell lines | eQTL | 8417 |
| GRASP 2 Catalog | 18193047 | Cells - Lymphoblastoid cell lines | eQTL | 463 |
| GRASP 2 Catalog | 18478092 | Cells - Leukemia cells | eQTL | 6 |
| GRASP 2 Catalog | 18478092 | Cells - Normal peripheral leukocytes | eQTL | 6 |
| GRASP 2 Catalog | 19128478 | Blood cells in celiac disease | eQTL | 2812 |
| GRASP 2 Catalog | 19222302 | Brain - Cortex | eQTL | 1281 |
| GRASP 2 Catalog | 19222302 | Cells - Peripheral blood mononuclear cells | eQTL | 2420 |
| GRASP 2 Catalog | 19361613 | Brain - Cortex with alzheimer's interaction | eQTL | 252 |
| GRASP 2 Catalog | 19361613 | Brain - Cortex with no alzheimer's interaction | eQTL | 1117 |
| GRASP 2 Catalog | 19644074 | Cells - Fibroblasts | eQTL | 512 |
| GRASP 2 Catalog | 19644074 | Cells - Lymphoblastoid cell lines | eQTL | 551 |
| GRASP 2 Catalog | 19644074 | Cells - T cells | eQTL | 532 |
| GRASP 2 Catalog | 19654370 | Osteoblasts | eQTL | 711 |
| GRASP 2 Catalog | 19680542 | Cells - Lymphoblastoid cell lines | eQTL | 73 |
| GRASP 2 Catalog | 19680542 | Osteoblasts | eQTL | 133 |
| GRASP 2 Catalog | 19966804 | Cells - Peripheral leukocytes | eQTL | 215 |
| GRASP 2 Catalog | 20084173 | Liver | eQTL | 49 |
| GRASP 2 Catalog | 20170901 | Cells - Aortic endothelial cells | eQTL | 5 |
| GRASP 2 Catalog | 20170901 | Cells - Aortic endothelial cells treated with OX-PAPC | eQTL | 35 |
| GRASP 2 Catalog | 20351726 | Brain - Prefrontal cortex | eQTL | 3035 |
| GRASP 2 Catalog | 20833654 | Cells - CD4+ lymphocytes | eQTL | 6843 |
| GRASP 2 Catalog | 20844574 | Blood | eQTL | 9 |
| GRASP 2 Catalog | 21226949 | Endometrial tumors | eQTL | 229 |
| GRASP 2 Catalog | 21637794 | Liver | eQTL | 7970 |
| GRASP 2 Catalog | 21789213 | Cells - Lymphoblastoid cell lines | eQTL | 8 |
| GRASP 2 Catalog | 21829388 | Adipose - Subcutaneous (Abdominal) | eQTL | 3 |
| GRASP 2 Catalog | 21829388 | Adipose - Visceral (Omentum majus) | eQTL | 6 |
| GRASP 2 Catalog | 21829388 | Blood | eQTL | 86912 |
| GRASP 2 Catalog | 21829388 | Musculus rectus - Abdominis | eQTL | 3 |
| GRASP 2 Catalog | 21949713 | Sputum | eQTL | 2047 |
| GRASP 2 Catalog | 22006096 | Liver | eQTL | 1376 |
| GRASP 2 Catalog | 22233810 | Cells - Dendritic cells | eQTL | 9434 |
| GRASP 2 Catalog | 22233810 | Cells - Dendritic cells treated with mycobacterium tuberculosis | eQTL | 9488 |
| GRASP 2 Catalog | 22383892 | Adipose - Gluteal | eQTL | 11 |
| GRASP 2 Catalog | 22383892 | Adipose - Subcutaneous (Abdominal) | eQTL | 30 |
| GRASP 2 Catalog | 22447449 | Liver and corresponding expression modules | eQTL | 28 |
| GRASP 2 Catalog | 22447449 | Liver | eQTL | 1 |
| GRASP 2 Catalog | 22509407 | Whole blood in amyotrophic lateral sclerosis cases and controls | eQTL | 15 |

| | | | | |
|---|---|---|---|---|
| GRASP 2 Catalog | 22522925 | Breast tumors | eQTL | 2981 |
| GRASP 2 Catalog | 22685416 | Brain - Cerebellum in alzheimer's disease cases and controls | eQTL | 20554 |
| GRASP 2 Catalog | 22685416 | Brain - Cerebellum in alzheimer's disease cases | eQTL | 3541 |
| GRASP 2 Catalog | 22685416 | Brain - Cerebellum in non-alzheimer's disease samples | eQTL | 3107 |
| GRASP 2 Catalog | 22685416 | Brain - Cerebellum in progressive supranuclear palsy cases | eQTL | 1610 |
| GRASP 2 Catalog | 22685416 | Brain - Temporal cortex in alzheimer's disease cases and controls | eQTL | 20069 |
| GRASP 2 Catalog | 22685416 | Brain - Temporal cortex in alzheimer's disease cases | eQTL | 3204 |
| GRASP 2 Catalog | 22685416 | Brain - Temporal cortex in non-alzheimer's disease samples | eQTL | 1747 |
| GRASP 2 Catalog | 22685416 | Brain - Temporal cortex in progressive supranuclear palsy cases | eQTL | 1117 |
| GRASP 2 Catalog | 22692066 | Whole blood | eQTL | 1470 |
| GRASP 2 Catalog | 22912676 | Cells - B-lymphoblastoid cell lines | eQTL | 25 |
| GRASP 2 Catalog | 23474282 | Ileum - Pre-pouch | eQTL | 71037 |
| GRASP 2 Catalog | 23671422 | Cells - Fibroblasts | eQTL | 1 |
| GRASP 2 Catalog | 23671422 | Cells - Fibroblasts, T cells and B-lymphoblastoid cell lines | eQTL | 2 |
| GTEx v7 | 23715323 | Adipose - Subcutaneous | eQTL | 1492748 |
| GTEx v7 | 23715323 | Adipose - Visceral (Omentum) | eQTL | 989804 |
| GTEx v7 | 23715323 | Adrenal gland | eQTL | 521055 |
| GTEx v7 | 23715323 | Artery - Aorta | eQTL | 1034149 |
| GTEx v7 | 23715323 | Artery - Coronary | eQTL | 338471 |
| GTEx v7 | 23715323 | Artery - Tibial | eQTL | 1547788 |
| GTEx v7 | 23715323 | Brain - Amygdala | eQTL | 150681 |
| GTEx v7 | 23715323 | Brain - Anterior cingulate cortex (BA24) | eQTL | 276912 |
| GTEx v7 | 23715323 | Brain - Caudate (Basal ganglia) | eQTL | 414047 |
| GTEx v7 | 23715323 | Brain - Cerebellar hemisphere | eQTL | 543823 |
| GTEx v7 | 23715323 | Brain - Cerebellum | eQTL | 780268 |
| GTEx v7 | 23715323 | Brain - Cortex | eQTL | 455391 |
| GTEx v7 | 23715323 | Brain - Frontal cortex (BA9) | eQTL | 327068 |
| GTEx v7 | 23715323 | Brain - Hippocampus | eQTL | 211589 |
| GTEx v7 | 23715323 | Brain - Hypothalamus | eQTL | 214764 |
| GTEx v7 | 23715323 | Brain - Nucleus accumbens (Basal ganglia) | eQTL | 357772 |
| GTEx v7 | 23715323 | Brain - Putamen (Basal ganglia) | eQTL | 278404 |
| GTEx v7 | 23715323 | Brain - Spinal cord (Cervical c-1) | eQTL | 167028 |
| GTEx v7 | 23715323 | Brain - Substantia nigra | eQTL | 112251 |
| GTEx v7 | 23715323 | Breast - Mammary tissue | eQTL | 695994 |
| GTEx v7 | 23715323 | Cells - EBV-transformed lymphocytes | eQTL | 271398 |
| GTEx v7 | 23715323 | Cells - Transformed fibroblasts | eQTL | 1278165 |
| GTEx v7 | 23715323 | Colon - Sigmoid | eQTL | 661130 |
| GTEx v7 | 23715323 | Colon - Transverse | eQTL | 784629 |
| GTEx v7 | 23715323 | Esophagus - Gastroesophageal junction | eQTL | 697016 |
| GTEx v7 | 23715323 | Esophagus - Mucosa | eQTL | 1474044 |
| GTEx v7 | 23715323 | Esophagus - Muscularis | eQTL | 1384504 |
| GTEx v7 | 23715323 | Heart - Atrial appendage | eQTL | 841069 |
| GTEx v7 | 23715323 | Heart - Left ventricle | eQTL | 740015 |
| GTEx v7 | 23715323 | Liver | eQTL | 305452 |
| GTEx v7 | 23715323 | Lung | eQTL | 1385916 |
| GTEx v7 | 23715323 | Minor salivary gland | eQTL | 125852 |
| GTEx v7 | 23715323 | Muscle skeletal | eQTL | 1438236 |
| GTEx v7 | 23715323 | Nerve - Tibial | eQTL | 1837615 |
| GTEx v7 | 23715323 | Ovary | eQTL | 262592 |
| GTEx v7 | 23715323 | Pancreas | eQTL | 696296 |
| GTEx v7 | 23715323 | Pituitary | eQTL | 552811 |
| GTEx v7 | 23715323 | Prostate | eQTL | 283528 |
| GTEx v7 | 23715323 | Skin - Not sun exposed (Suprapubic) | eQTL | 1250843 |
| GTEx v7 | 23715323 | Skin - Sun exposed (Lower leg) | eQTL | 1697927 |
| GTEx v7 | 23715323 | Small intestine - Terminal ileum | eQTL | 302654 |
| GTEx v7 | 23715323 | Spleen | eQTL | 494717 |
| GTEx v7 | 23715323 | Stomach | eQTL | 596021 |
| GTEx v7 | 23715323 | Testis | eQTL | 1417129 |
| GTEx v7 | 23715323 | Thyroid | eQTL | 2017804 |
| GTEx v7 | 23715323 | Uterus | eQTL | 169796 |
| GTEx v7 | 23715323 | Vagina | eQTL | 167241 |
| GTEx v7 | 23715323 | Whole blood | eQTL | 1001052 |
| Haploreg v4.1 | 17873874 | Cells - Lymphoblastoid cell lines | eQTL | 9979 |
| Haploreg v4.1 | 18462017 | Liver | eQTL | 3603 |
| Haploreg v4.1 | 20220756 | Cells - Lymphoblastoid cell lines | eQTL | 2049 |
| Haploreg v4.1 | 20485568 | Brain - Cerebellum | eQTL | 5233 |
| Haploreg v4.1 | 20485568 | Brain - Frontal cortex | eQTL | 5422 |
| Haploreg v4.1 | 20485568 | Brain - Pons | eQTL | 3377 |
| Haploreg v4.1 | 20485568 | Brain - Temporal cortex | eQTL | 5204 |
| Haploreg v4.1 | 21283786 | Osteoblasts BMP2 | eQTL | 1533 |

| | | | | |
|---|---|---|---|---|
| Haploreg v4.1 | 21283786 | Osteoblasts DEX | eQTL | 1248 |
| Haploreg v4.1 | 21283786 | Osteoblasts | eQTL | 1089 |
| Haploreg v4.1 | 21283786 | Osteoblasts PGE2 | eQTL | 1519 |
| Haploreg v4.1 | 23209423 | Lung | eQTL | 14977 |
| Haploreg v4.1 | 24604202 | Cells - Monocytes | eQTL | 5982 |
| Haploreg v4.1 | 24604202 | Cells - Monocytes IFN | eQTL | 9570 |
| Haploreg v4.1 | 24604202 | Cells - Monocytes LPS24 | eQTL | 2458 |
| Haploreg v4.1 | 24604202 | Cells - Monocytes LPS2 | eQTL | 3864 |
| Haploreg v4.1 | 24846176 | Heart | eQTL | 5627 |
| Haploreg v4.1 | 25174004 | Brain - Aveall | eQTL | 10597 |
| Haploreg v4.1 | 25174004 | Brain - Cerebellum | eQTL | 2094 |
| Haploreg v4.1 | 25174004 | Brain - Frontal cortex | eQTL | 1200 |
| Haploreg v4.1 | 25174004 | Brain - Hippocampus | eQTL | 1003 |
| Haploreg v4.1 | 25174004 | Brain - Inferior olivary nucleus | eQTL | 831 |
| Haploreg v4.1 | 25174004 | Brain - Intralobular white matter | eQTL | 1515 |
| Haploreg v4.1 | 25174004 | Brain - Occipital cortex | eQTL | 954 |
| Haploreg v4.1 | 25174004 | Brain - Putamen | eQTL | 550 |
| Haploreg v4.1 | 25174004 | Brain - Substantia nigra | eQTL | 415 |
| Haploreg v4.1 | 25174004 | Brain - Temporal Cortex | eQTL | 1391 |
| Haploreg v4.1 | 25174004 | Brain - Thalamus | eQTL | 841 |
| Kim et al. (Nature Comm 2017) | 28814792 | Cells - Monocytes | eQTL | 2403 |
| Kim et al. (Nature Comm 2017) | 28814792 | Cells - Monocytes (stimulated with LPS/IVT/MDP vs. unstimulated) | reQTL | 369 |
| pGWAS | 28240269 | Blood plasma | pQTL | 508 |
| ScanDB | 19933162 | Brain - Cerebellum | eQTL | 621247 |
| ScanDB | 19933162 | Brain - Parietal lobe | eQTL | 592490 |
| ScanDB | 19933162 | Liver | eQTL | 285990 |
| seeQTL | 17982457 | Brain | eQTL | 39154 |
| seeQTL | 22171328 | Cells - Lymphoblastoid cell lines | eQTL | 123727 |
| The Cardiogenics Project | 23382694 | Cells - Monocytes | eQTL | 1329875 |
| The Cardiogenics Project | 27558669 | Cells - Macrophages | eQTL | 1032604 |
| Zeller et al. (PlosONE 2010) | 20502693 | Cells - Monocytes | eQTL | 203642 |

**Supplementary Table 2.** For each of the investigated LD thresholds ($r^2$) top 100 genes with highest number of datasets per eQTL (n_max), annotated by mean expression percentile in GTEx (mean_expr_percentile) and by housekeeping gene status (is_housekeeping). Table is available in attached Excel sheet.

**Supplementary Table 3.** For each LD threshold the number of housekeeping genes and the number of genes in the 10% highest expressed genes in GTEx v8 among the 100 genes with the highest dataset count per eQTL LD block. Adjusted (FDR < 0.05) P-values were calculated using hypergeometric test assuming an overall gene number of 18,521 (genes with at least one best significant eQTL), a total of 3,705 housekeeping genes and a total of 1,201 genes among the 10% highest expressed genes.

| **LD threshold** | **#Housekeeping** | **Adjusted P** | **90th percentile expression** | **Adjusted P** |
|---|---|---|---|---|
| No LD | 29 | 1.84E-02 | 9 | 0.20 |
| r2 = 1 | 33 | 1.65E-03 | 12 | 0.06 |
| r2 > 0.95 | 42 | 2.12E-06 | 14 | 0.03 |
| r2 > 0.9 | 41 | 3.12E-06 | 12 | 0.06 |
| r2 > 0.8 | 37 | 1.05E-04 | 10 | 0.17 |
| r2 > 0.6 | 36 | 1.89E-04 | 9 | 0.20 |

**Supplementary Table 4.** Members of the Cardiogenics Consortium.

| | |
|---|---|
| Tony Attwood | Seraya Maouche |
| Stephanie Belz | Jasbir S Moore |
| Peter Braund | Gilles Montalescot |
| Jessy Brocheton | David Muir |
| François Cambien | Elizabeth Murray |
| Jason Cooper | Chris P Nelson |
| Abi Crisp-Hihn | Jessica Neudert |
| Panos Deloukas | David Niblett |
| Patrick Diemert | Karen O'Leary |
| Jeannette Erdmann | Willem H. Ouwehand |
| Nicola Foad | Helen Pollard |
| Tiphaine Godefroy | Carole Proust |
| Alison H. Goodall | Angela Rankin |
| Jay Gracey | Augusto Rendon |
| Emma Gray | Catherine M Rice |
| Rhian Gwilliams | Hendrik Sager |
| Susanne Heimerl | Nilesh J. Samani |
| Christian Hengstenberg | Jennifer Sambrook |
| Jennifer Jolley | Gerd Schmitz |
| Unni Krishnan | Michael Scholz |
| Heather Lloyd-Jones | Laura Schroeder |
| Ulrika Liljedahl | Heribert Schunkert |
| Ingrid Lugauer | Jonathan Stephens |
| Per Lundmark | Ann-Christine Syvannen |
| Chris Wallace | Stefanie Tennstedt |
